# Mapping cross-domain drivers of Alzheimer’s disease risk through integrated network analysis

**DOI:** 10.1101/2025.10.16.682921

**Authors:** Gregory A Cary, Stephen Keegan, Jesse C Wiley, Jake Gockley, Frank M Longo, Allan I Levey, Anna K Greenwood, Karina Leal, Gregory W Carter, The Emory-Sage-SGC-JAX TREAT-AD Center

**Affiliations:** The Jackson Laboratory, Bar Harbor, ME USA; University of Kansas, Lawrence, KS USA; Sage Bionetworks, Seattle, WA USA; Stanford University School of Medicine, Stanford, California USA; Emory University School of Medicine, Atlanta, GA USA

**Author notes:** Corresponding author: Gregory W Carter, The Jackson Laboratory, Bar Harbor ME. adknowledgeportal.org/TREATAD-ESS-Center. Abbreviations: Alzheimer’s Disease (AD), Target Enablement to Accelerate Therapy Development in AD (TREAT-AD), Accelerating Medical Partnership for AD (AMP-AD), Gene Set Enrichment Analysis (GSEA), Gene Ontology (GO), weighted Key Driver Analysis (wKDA), Target Enabling Package (TEP).

**Keywords:** Alzheimer’s Disease, Biological Domain, Network, Key Driver

## Abstract

**Introduction:** Alzheimer’s Disease (AD) is a complex neurodegenerative disorder with numerous known risk factors. Identification of which genetic factors are causal drivers is difficult due to the long disease prodrome in an inaccessible organ. The application of integrative, systems-level approaches are crucial for addressing this complexity.

**Methods:** Sixteen biological domain specific interaction networks were derived from the top AD risk-enriched proteins within each domain. Weighted key driver analysis identified influential hub nodes within each network.

**Results:** Distinct processes and drivers were identified within each domain’s network. Domains including Structural Stabilization, Endolysosome, and Lipid Metabolism were especially influential. Integrating key drivers across domains identified consistent drivers such as CTNNB1, ACSL1, and ALDH3A2, suggesting fundamental roles contributing to AD risk.

**Discussion:** This highly integrative network-based approach identified context-dependent drivers and enabled the inference of interactions between domains. The identified drivers suggest potential targets for future therapeutic development.

## 1. BACKGROUND

Alzheimer’s disease (AD) is a complex neurodegenerative disease with numerous genetic and environmental risk factors. While genome wide association studies implicate over 75 genomic loci in disease risk [1] and omics analyses have nominated over 900 candidate targets (agora.adknowledgeportal.org), how this risk translates to neurodegenerative phenotypes remains unclear. AD typically manifests late in life, after an extended prodromal period marked by neuropathological accumulation [2]. The intricate interplay between risk-implicated biological processes over decades in an inaccessible organ makes identifying causal genes driving AD progression extremely difficult. This ambiguity then hinders the development of disease modifying therapies, which are desperately needed given the increasing prevalence and socioeconomic and health system burdens imposed by AD [3].

Our previous work established an integrated risk scoring framework to summarize genetic and multi-omic risk, and defined AD biological domains (biodomains) representing commonly implicated molecular processes [4]. These resources allow partitioning of disease risk signatures into distinct, more analyzable components for focused inquiry into pathophysiology. While biodomains provide valuable insights into disease-associated endophenotypes, they are still fundamentally lists of genes with similar function, which fail to capture the intricacies of disease mechanisms. In contrast, a network-based approach considers the relationships between the genes underlying specific biological processes, leveraging systems-level knowledge to identify influential hub genes with causal roles in the system as a whole and its degradation in AD.

Of these approaches, molecular interaction network graphs are ideal for representing biological entities (e.g. genes or proteins) as nodes and relationships between entities (e.g. pathways or biochemical interactions) as edges. Similar approaches have been used to prioritize risk associated genes [5] and identify clinically successful therapeutic targets [6]. The success of network-based approaches is due to the observation that proteins that are closely connected within biological networks often share functional roles and impact similar phenotypes, including disease [7,8]. Another principle that emerges from these efforts is that comprehensive networks that comprise data from multiple sources tend to out-perform smaller networks in the identification of disease genes [9,10].

Network analyses have also been successfully applied to AD data — including to identify regulators of selective neuronal vulnerability to AD neurodegeneration [11,12], as prior knowledge for graph neural network based machine learning approaches [13], and to elucidate key drivers of multi-scale expression networks [14]. There is tremendous potential for biodomain-specific networks to identify context-dependent drivers of AD.

This paper utilizes TREAT-AD derived genetic and multi-omic risk in the generation of network graphs germane to each AD-relevant biodomain and subsequently identifies influential hub nodes, or key driver genes. These key drivers represent pivotal regulatory proteins for biological processes represented within each graph; processes which frequently span other biodomains. We start by performing a shortest path reconstruction between all risk-enriched proteins from each biodomain across the composite Pathway Commons network graph. The weighted graph reconstruction is dictated by both disease risk scores [4] and annotation to the target biodomain, which results in 16 network graphs that are biodomain-specific. Next, weighted key driver analysis (wKDA) identifies proteins whose variation influences sets of genes annotated to biodomain-specific terms. The drivers identified show distinct sets of target processes depending on which graph is analyzed. For example, nodes in the Endolysosome graph are primarily drivers for Synapse domain processes, suggesting that impaired endolysosomal trafficking may drive synaptic dysfunction in AD by disrupting protein turnover and receptor recycling. We use such inferences to build a network of domain interactions that identifies the Structural Stabilization and Endolysosome graphs as being among the most influential domains. Finally, we integrate wKDA results across multiple domain contexts and show that the key drivers that are consistently identified represent fundamental drivers of AD risk.

## 2. METHODS

### 2.1 Interaction graph data and annotation

We used the Pathway Commons database v12 [15] as a basis for generating biodomain focused interaction graphs. This is a comprehensive database consisting of interactions captured from 18 other data sources, including pathway databases (e.g. KEGG and Reactome) as well as protein-protein interaction databases (e.g. BioGRID and IntAct) (Figure S1A). In total this resource captures 1.2 million interactions among over 19 thousand proteins (Figure 1B-C).

**Figure 1.**
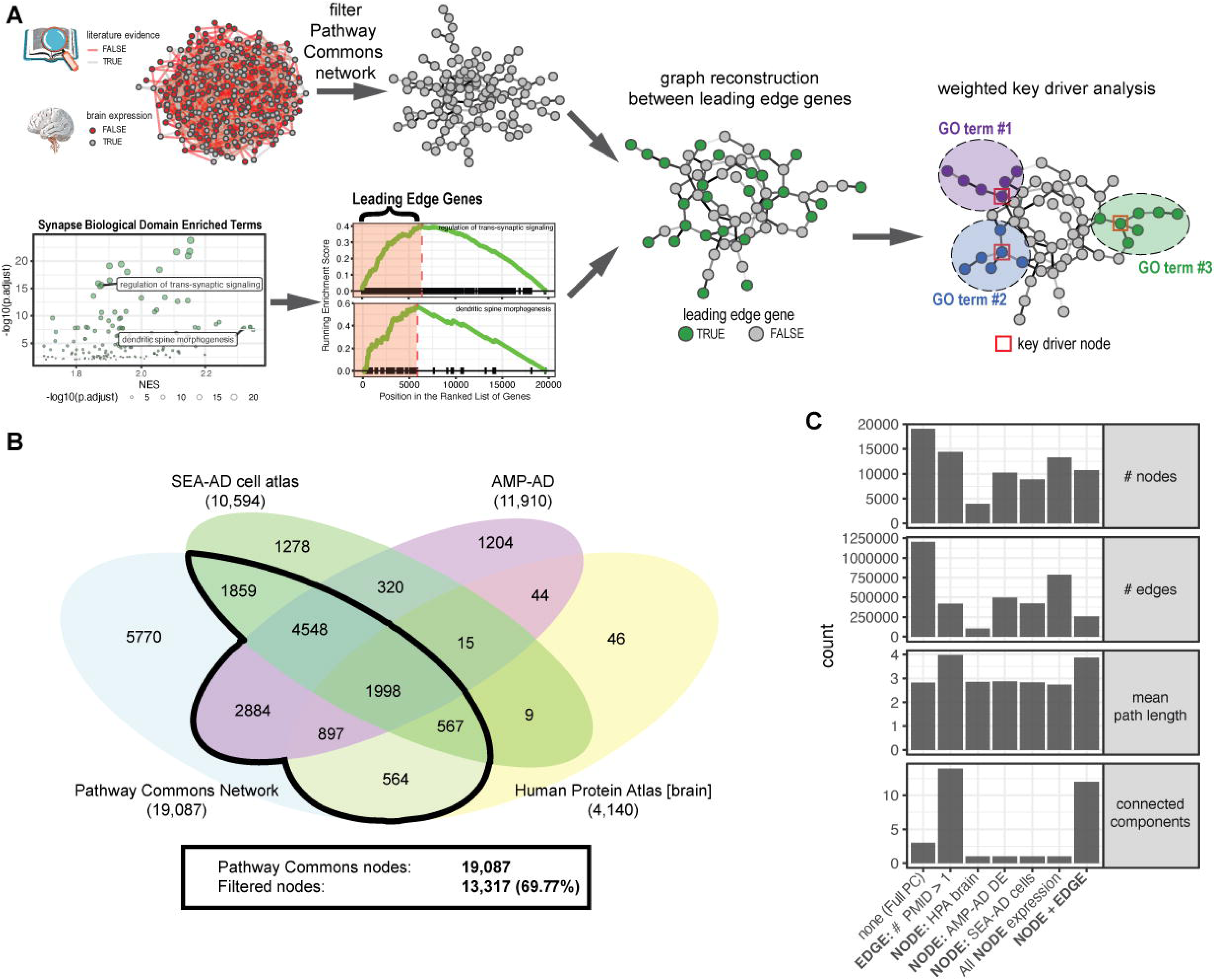
Methodological overview and Pathway Commons network filtering. (A) Overview of the methodology employed to generate biodomain specific networks and identify key driver hub nodes. The Pathway Commons network was filtered based on literature evidence and brain expression (top), and leading edge genes from enriched biodomain terms (here for the Synapse domain) were used to seed biodomain network generation. Finally, weighted key driver analysis was performed on each biodomain graph to identify influential hub nodes. (B) Venn diagram showing the Pathway Commons network nodes with support for brain expression from the Human Protein Atlas, AMP-AD differential expression analysis, and SEA-AD single cell analysis workflows. (C) Network statistics under various filtering conditions; the final category (“NODE + EDGE”) was used for biodomain network generation.

The edges in the Pathway Commons network are annotated with supporting literature where available (Figure S1A). In addition, we annotated each node for evidence of expression in the brain using any three different sources. We examined evidence of expression in normal brain tissues using the Human Protein Atlas (HPA) tissue microarray data v22 [16]. We assembled the list of proteins for which there was evidence (Reliability either “Enhanced”, “Supported”, or “Approved”) for any level of expression (Level either “Low”, “Medium”, “High”, “Ascending”, or “Descending”) for any of the nine brain tissues included in the normal tissue IHC data (i.e. "caudate", "cerebellum", "cerebral cortex", "hippocampus", "hypothalamus", "pituitary gland", "dorsal raphe", "choroid plexus", or "substantia nigra"). Using these criteria we identified 4,140 proteins with plausible brain expression in normal human brain tissue. We also assembled a list of proteins for which there was reliable evidence of no expression (i.e. Level is “Not Detected”) in any of the brain tissues sampled (N = 1,324 proteins) to benchmark our filtering.

We also included genes and proteins for which we have reliable evidence of differential expression in AD versus control brains. Significant transcriptional or proteomic differential expression (FDR ≤ 0.05) from our meta-analysis of AMP-AD datasets [4] yielded a set of 11,910 genes. We then compared this expression with the RNA GTEx brain region data from the HPA dataset above (Figure S1B). The normalized TPM distribution for genes identified as reliably brain expressed in the HPA tissue IHC data and those that are significantly differentially expressed in AD brains was similarly distributed, and consistently higher than for those genes reliably not expressed in the brain using the HPA data.

Finally, recognizing that some genes may not be strongly expressed in bulk tissue, we examined evidence of expression in brain cell types using data from the Seattle Alzheimer’s Disease Brain Cell Atlas (SEA-AD) [17]. We computed a 90^th^ percentile expression and the fraction of cells expressing each gene in each Supertype. We identified a threshold that separated genes that were either reliably brain expressed or differentially expressed in AMP-AD datasets from those genes that were reliably not expressed in brain tissues (Figure S1C). When we applied this filter we removed 89% of the genes that are not expressed in the brain while retaining 60% of AMP-AD differentially expressed genes and 64% of the HPA brain expressed genes (Figure S1D).

Each node in the Pathway Commons graph was annotated with a binary label as to whether it has evidence of expression in the brain from each of these sources. The nodes were also annotated to include the TREAT-AD Target Risk Scores [4]. The fully annotated base network graph is available through Synapse (syn51110930). The starting graph for each biodomain specific graph was this base graph filtered to include nodes with any evidence of expression in the brain and edges with at least two distinct PubMed IDs supporting their existence.

### 2.2 Dijkstra shortest path reconstruction of biodomain graphs

To seed the generation of biodomain specific network graphs, we used the results of gene set enrichment analysis (GSEA) using the AD Target Risk Score as a ranking statistic. We used the gseGO function from the clusterProfiler R package (v4.6.2) [18,19] using the org.Hs.eg.db annotation database (v3.16) and the resulting gene ontology terms were categorized into biodomains based on the GO ID of enriched terms ([4], syn25428992). To focus the graph construction around genes from the top of the risk score distribution for each domain, we selected terms that were significant in this analysis with a Benjamini-Hochberg adjusted p-value ≤ 0.01 and a normalized enrichment score above 1.66 (Figure S2A). The leading edge genes from all terms in a domain meeting these criteria were then used as the input list to generate biodomain network graphs.

For each gene in the input list, we used Dijkstra’s algorithm to identify the shortest path in the filtered base network between that gene and all others in the input list. Dijkstra’s algorithm relies on edge weights to compute the shortest distance between two nodes. The edge weights for the graph reconstruction were set to be the sum of 5 (i.e. the maximum possible Target Risk Score) minus the Target Risk Score for each node joined by the edge, and then an additional weight for whether both nodes were annotated to the target domain (0), if only one node was annotated to the target domain (3), or if neither node was annotated to the target domain (5). Thus the largest possible weight for each edge is 15, and the smallest edge weights (i.e. shortest distances for Dijkstra’s algorithm) were focused on the nodes with the largest Target Risk Score and those that were annotated to the target domain. The final network graph for each biodomain is accessible on Synapse (syn69925494). We tested for over-represented terms among the biodomain graphs using the enrichGO function from the clusterProfiler R package [18,19] against the org.Hs.eg.db annotation database (v3.16) and categorized significantly enriched terms into biodomains based on the term ID.

### 2.3 Weighted key driver analysis

We used the wKDA module from the Mergeomics R package ([20,21], Mergeomics_Version_1.99.0.R, github.com/jessicading/mergeomics). This package takes a network file, formatted as an edge list where each edge is assigned a weight, and a module file with groups of nodes around which the key driver nodes should be identified. For the wKDA analyses the edge weights were simply the sum of the Target Risk Scores for each node connected by the edge. For the module file, we used the list of all genes annotated to GO terms within the biodomain framework. The wKDA was run with the following parameters: search depth was set to 1, number of permutations was set to 2000, minimum module size was set to 5, and the networks were analyzed as undirected.

Redundant edges between nodes of different types or from different classes were simplified to only retain one edge per interaction. For each network we ran wKDA twice with different edge factor parameter settings; both with the edge factor parameter set to 1, for which wKDA considered the AD risk score edge weighting in the analyses of hub nodes, and with the edge factor set to 0 for which wKDA treated all edges as having uniform weights. We then joined the results of the unweighted (edge factor 0) and AD risk weighted (edge factor 1) analyses within the same network by the same nodes and the same GO term module and computed a ΔFDR as the difference between the -log10 transformed FDR values between the weighted and the unweighted analyses. This difference represents a topologically normalized, directed metric of AD risk influence between proteins. We noted that in the analyses some predicted drivers were asserted for GO terms that had few proteins present in the network. We excluded from our key driver list any driver-term pair for which fewer than 25% of the term proteins were present in the network.

### 2.4 Integrating key drivers across graphs

To identify the most consistent pairings of key drivers with biodomain terms across biodomain networks we summarized the key driver results across networks. We only considered drivers that were identified with a risk weighted FDR (i.e. edge factor 1 analysis) less than or equal to 0.05 and where at least 25% of the proteins annotated to the term were represented in the network. The wKDA statistics for each driver were dependent on the size of the originating graph, and so for each graph we ranked the risk weighted (i.e. edge factor 1 analysis) -log10 FDR as well as the ΔFDR of every pair of key driver nodes with GO terms. We scaled down the ranked FDR and ΔFDR values for any driver that was identified as a candidate, but not the top driver for the term in the graph, by multiplying the ranks by 0.33. In this way the top driver for each term in the graph has the highest rank, but if a node is identified as a candidate hub it is still considered in the integrative analysis in case it is a key driver in other networks as well. Finally, we multiplied the scaled, ranked FDR and ΔFDR values by the proportion of the GO term present in the graph to emphasize the predicted drivers for processes that are well represented in the graph. For each pairing of key driver and GO term, we then summed the ranked KDA statistics across all graphs to identify the most consistently strong drivers across networks.

### 2.5 Additional data and code

In our analyses of our driver sets we compare with other gene sets. These include the set of key drivers from another analysis of AD co-expression based Bayesian networks [14]. These key drivers were identified at multiple scales (i.e. transcriptome, proteome, and global). For our analyses we considered a union of all identified drivers across the scales in the Beckmann study (N = 702). We also considered the list of all currently nominated targets from the AMP AD consortium, as cataloged on the Agora site (agora.adknowledgeportal.org/genes/nominated targets) (N = 810). The set of AD GWAS genes (N = 123) was derived by taking the union of lists from three sources: the supplementary table that accompanies Neuner *et al* [22] (N = 71), supplementary table S5 from Bellenguez *et al* [1] that identifies all genome wide significant loci (N = 77), and the list of AD loci with genetic evidence compiled by the ADSP Gene Verification Committee (adsp.niagads.org/gvc-top-hits-list) (N = 76).

We also considered the known AD cell type expression and phenotypic effects of knockdown for identified key drivers, in particular ALDH3A2. For phenotypic effects we referred to the CRISPRbrain.org resource [23] and focused on the genome wide CRISPRa screen of glutamatergic neuron survival following activation of genes. Cell type expression in AD was assessed using data available from SEA-AD [17]. Specifically we queried the transcriptomic effects across disease pseudoprogression for cell Subclass using the available shiny application (sea- ad.shinyapps.io/ad_gene_trajectories).

All biodomain graphs and key driver analysis results generated through this work are available on Synapse (syn69925494). All code to reproduce these analyses has been made available in the GitHub repository (github.com/caryga/treatad_biodomain_graph_kda_paper).

## 3. RESULTS

The overall analytical strategy employed in this work is depicted (Figure 1A). This includes annotating and filtering the base network graph from the Pathway Commons database to only include nodes with evidence of expression in the brain and edges with multiple publications supporting the existence of a molecular interaction. For each biodomain we identify a set of input nodes that are the most risk-enriched genes (i.e. leading edge genes) from all significantly enriched domain terms based on gene set enrichment analyses. The weighted reconstruction around these leading edge genes using the filtered Pathway Commons network generates domain-specific graphs with reduced network redundancy while allowing for the inclusion of important interactors that were not in the leading edge gene set. Influential hub nodes are then identified in each domain graph using a weighted key driver analysis.

### 3.1 Biodomain Graph Construction

To generate domain-specific knowledge graphs we leveraged the Pathway Commons database [15]. In the Pathway Commons network, each edge is annotated with PubMed IDs of references supporting its existence, where available. We supplemented this by annotating each node for evidence of expression in the brain using any of: (1) the Human Protein Atlas tissue microarray data [16], (2) evidence of differential expression in AD brains from our meta-analysis of AMP-AD datasets [4], or (3) evidence of expression in a brain cell type using data from the Seattle Alzheimer’s Disease Brain Cell Atlas (SEA-AD) [17] (Methods, Figure S1B-D). Of the 19,087 nodes in the Pathway Commons knowledge graph, there was evidence for brain expression for 13,317 (69.77%) (Figure 1B). Out of all 16,233 genes for which there was evidence for brain expression, 2,916 (17.96%) were not found in the Pathway Commons graph, and these genes tended to have a lower AD Target Risk Score (Figure S1E, Kolmogorov-Smirnov p-value = 0.0005). When we filtered the Pathway Commons graph to only include nodes with evidence of brain expression and edges with more than one PubMed ID supporting evidence, we were left with a graph consisting of 10,722 nodes (56.2%) and 256,850 edges (21.3%), which is the network used as the basis for the reconstruction of the biodomain graphs (Figure 1C).

Each biodomain graph was constructed around a set of genes from significantly enriched GO terms from that domain. We performed gene set enrichment analysis using the AD Target Risk Score and annotated all enriched terms to the AD biodomains [4]. We included terms that were significant with an adjusted p-value ≤ 0.01 and a normalized enrichment score above 1.66 (Figure S2A). Using these thresholds left no significantly enriched terms from three domains: RNA Spliceosome, DNA Repair, and Epigenetic. The remaining 574 significantly enriched terms are distributed non-uniformly across the 16 biodomain, with 165 significant terms from the Synapse domain and only 2 significant terms from the Tau Homeostasis domain (Figure S2B). The leading edge genes from all of the terms within a biodomain were used to seed the graph reconstruction for each domain. There is generally good correspondence between the number of significant terms and the number of leading edge genes, with Synapse having the largest number of leading edge genes (n = 1496) and Tau Homeostasis having the fewest (n = 37) (Figure S2B). The Synapse, Immune Response, and APP Metabolism domains have the fewest leading edge genes per GO term, while the Apoptosis and Proteostasis domains have the most leading edge genes per GO term (Figure S2B). Not all leading edge genes are present in the filtered base graph; across domains there are about 5-10% of leading edge genes that are absent from the base graph and therefore not able to be included in the final biodomain graph (Figure 2SB). The domains with the largest proportion of leading edge genes missing from the base graph include Myelination (13.4%) and Synapse (12.2%). The overlap between the nodes in the filtered base graph and the leading edge genes for a given domain was the input gene set for domain-specific graph generation.

**Figure 2.**
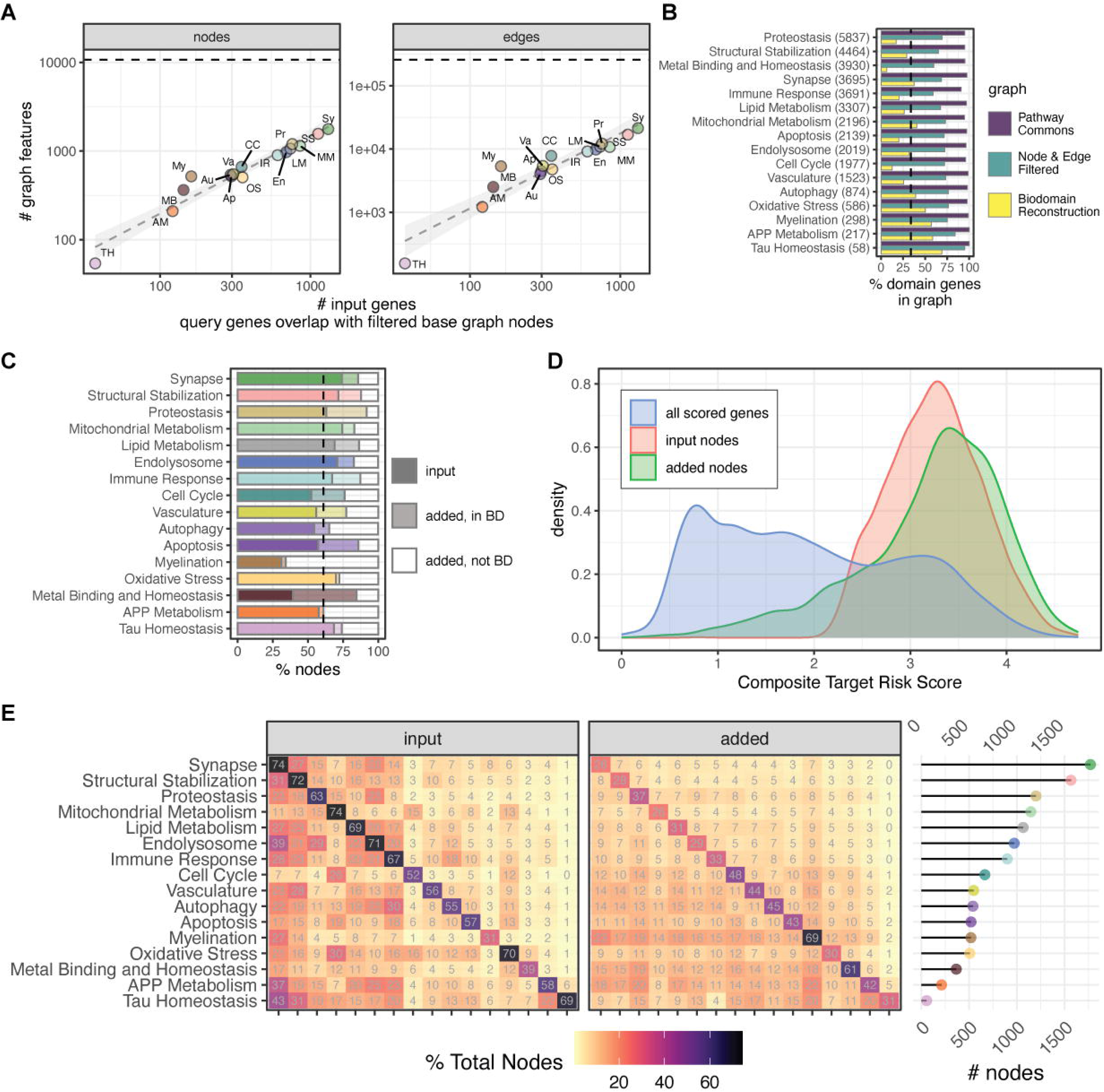
Properties of Biodomain-specific Networks. (A) The number of nodes and edges in each biodomain graph (y-axis, log scale) as a function of the number of input genes (x-axis, log scale). Each point represents a different biodomain: Tau Homeostasis (TH), APP Metabolism (AM), Metal Binding and Homeostasis (MB), Myelination (My), Autophagy (Au), Apoptosis (Ap), Oxidative Stress (OS), Vasculature (Va), Cell Cycle (CC), Immune Response (IR), Endolysosome (En), Lipid Metabolism (LM), Proteostasis (Pr), Mitochondrial Metabolism (MM), Structural Stabilization (SS), and Synapse (Sy). (B) Percentage of biodomain genes present within graphs across different stages of graph generation: the overall Pathway Commons graph, the node and edge filtered graph, and the final biodomain-specific graph. The total number of genes in each domain is shown parenthetically on the y-axis and the dashed line represents the average percent of domain genes present in the final graphs across domains. (C) Proportion of network nodes in final biodomain graphs that were used as input versus those that were added during reconstruction. Nodes that were added were further subdivided into those annotated to the domain and those that are not annotated to the domain. (D) Distribution of composite AD Target Risk Scores biodomain graph nodes, subdivided into input nodes and added nodes, as well as the background of all scored targets. (E) Overlap of nodes across biodomain graphs. The heatmaps show the percentage overlap of input and added nodes between biodomain graphs. The total number of nodes in each graph is shown on the right. The order of biodomains along the x-axis is the same as the order along the y-axis.

Since a goal of this effort was to focus graph construction around AD risk genes partitioned by biodomain, graph network reconstruction with Dijkstra’s algorithm calculated the shortest paths between each input node and all other input nodes for a given domain using edge weights, or distances between nodes.The edge weights used for construction were a combination of AD Target Risk Scores for the two nodes connected by an edge and a score for whether the nodes are annotated to the biodomain being targeted (see Methods for more details). This method of graph network reconstruction reduces edge redundancy and paths that are not relevant to AD, while allowing essential interacting nodes that were not leading edge genes in the biodomain to remain.

The number of nodes and edges in the resulting graphs was proportional to the number of input nodes (Figure 2A). The average graph size across all domains is 784 nodes and 7916 edges, with the largest graph belonging to the Synapse domain (1,770 nodes and 21,365 edges) and the smallest graph belonging to the Tau Homeostasis domain (54 nodes and 156 edges) (Figure S3A, Supplemental Table 1). The graphs show general signs for high global connectivity (i.e. low number of connected components and high average node degree), but this high connectivity is not arranged around nodes of similar degrees (assortativity coefficients near 0), suggesting the absence of clusters of high-degree nodes (Figure S3A). Considering the proportion of genes from each biodomain present in each graph class, there are over 90% of genes annotated to each biodomain present in the full Pathway Commons graph, between 59%-95% of biodomain genes present in the filtered base graph, and 6%-69% of biodomain genes present in each reconstructed graph (Figure 2B).

The shortest path reconstruction adds nodes to each graph. On average there were 275 nodes added, reflecting 38.9% (ranging 26% to 69%) of the total nodes in the resulting biodomain graphs that were added (Figure 2C and 2E). The majority of added nodes are one node away from an input node, though a few added nodes require longer paths, as many as 6 or 7 nodes not including the input nodes (Figure S3A, minimum path length of added nodes). The added nodes reflect a mixture of both nodes that are annotated to the target domain and those that are not, but presumably have higher AD risk scores. There is a broader distribution of risk scores for added nodes (Figure 2D).

Comparing nodes across biodomain graphs, only about 10%-20% of nodes added in each domain overlap with nodes added in other domains (Figure 2E), suggesting that the nodes added during reconstruction are generally domain-specific. While there is a slightly higher proportion of overlapping input nodes, there is a limited overlap of graph nodes between biodomains. The largest percentage of shared nodes tend to involve the smallest graphs (e.g. APP Metabolism, Tau Homeostasis, and Myelination), which all share over 70% of their nodes with the largest graph (i.e. Synapse). Another measure of the domain-specificity of each graph is from the examination of GO term over- representation analysis of nodes from each graph (Figure S3B). The most significantly overrepresented GO terms for each set of graph nodes corresponds to the target domain, which makes sense given that the input lists were derived from genes annotated to GO terms within that biodomain. However, there are also differences between the overrepresented terms from the domain graph nodes, excluding the target domain. For example, Autophagy nodes are strongly enriched for Immune Response and Endolysosome terms, while Cell Cycle nodes are strongly enriched for Mitochondrial Metabolism terms. These off-target domain enrichments imply potentially important functional intersections between domains.

### 3.2 Weighted Key Driver Analysis of Biodomain Graphs

To identify key drivers (i.e., influential hub nodes) within the biodomain graphs we used the weighted key driver analysis (wKDA) module from the Mergeomics R package [20,21]. This software scans each graph for hub nodes and assesses overrepresentation of hubs connected with specified gene sets. For the current study we used as gene sets the list of all biodomain GO terms and the genes annotated to each. The Mergeomics wKDA module allows for the control of how influential edge weights are in the identification of key drivers. For our purposes we performed wKDA on each biodomain graph twice, once with uniform edge weighting (i.e. edge factor 0) and once with edge weights based on the AD Target Risk Scores of the two connected nodes (i.e. edge factor 1). We then combined the results of each wKDA run, joining the statistics on both the identity of the driver node and the gene sets being driven, and compared the difference in the significance of each driver identified in the risk weighted analysis versus the uniform edge weight analysis (ΔFDR) (Figure S4A, Supplemental Table 2). This informs which drivers are identified based primarily on the topology of the examined sub-network versus those drivers that are most influenced by the inclusion of disease risk. Finally, while most biodomain GO terms have over 75% genes annotated to the term present in the full Pathway Commons and filtered base networks, many biodomain GO terms only have a fraction of annotated genes present in the reconstructed biodomain graphs (Figure S4B-D). The key driver results were filtered to only consider nodes as key drivers if the term being driven has over 25% of annotated genes represented in the graph.

Our analyses identified 701 key driver relationships involving 192 unique driver nodes across 422 unique GO terms (Table 1, Supplemental Table 2). In general the number of key drivers identified scales with the number of nodes in each graph, with larger graphs yielding more key drivers (Figure 3A). No drivers were identified in the Tau Homeostasis graph, likely because the graph is so sparse. Between 1.5% and 6% of all nodes in the biodomain graphs are identified as key drivers (Figure 3B), with the Endolysosome, Structural Stabilization, and Lipid Metabolism graphs having the largest proportions of nodes identified as key drivers. Many of the identified key drivers were also identified as key drivers in previous analysis of AD expression-based networks (N = 62, Fisher p = 3.9 x 10^-22^, OR = 5.4) [14] or have been nominated for follow-up by the AMP-AD consortium (N = 51, Fisher p = 9.5 x 10^-13^, OR = 3.7) [24,25]. Only four of the 276 top key drivers identified are also GWAS loci (*APP*, *INPP5D*, *NCK2*, *PRNP;* Fisher p = 0.16, OR = 1.98) and while this is not a significant overlap it is consistent with the observation that targets with genetic support are over twice as likely to be successful in clinical trials [26].

**Figure 3.**
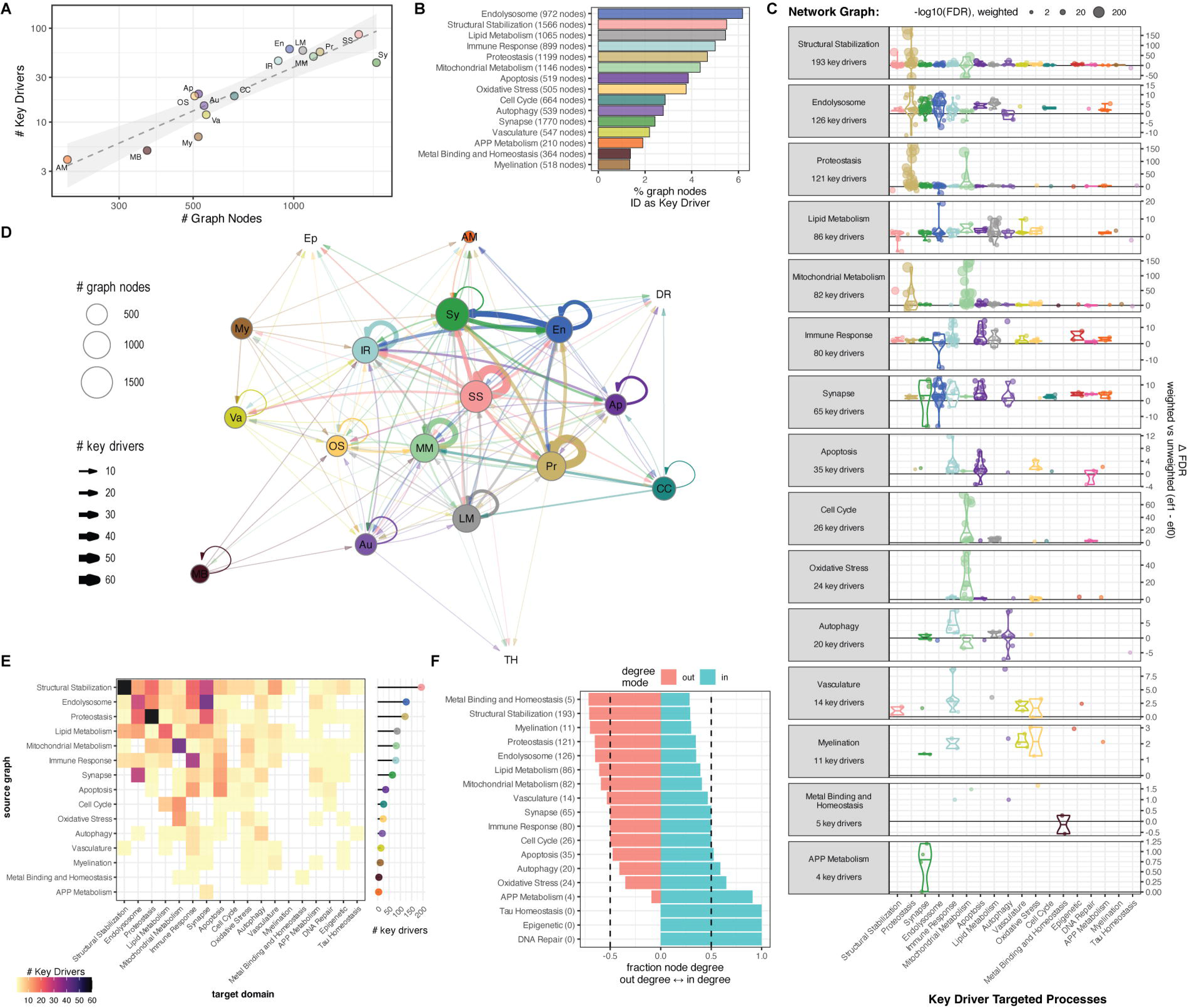
Key driver analysis of AD biodomain networks. (A) Scatter plot showing the number of key driver nodes identified within each biodomain graph (y-axis) versus the total number of nodes present in each graph (x-axis). Each point represents a different biodomain graph: APP Metabolism (AM), Metal Binding and Homeostasis (MB), Myelination (My), Autophagy (Au), Apoptosis (Ap), Oxidative Stress (OS), Vasculature (Va), Cell Cycle (CC), Immune Response (IR), Endolysosome (En), Lipid Metabolism (LM), Mitochondrial Metabolism (MM), Proteostasis (Pr), Structural Stabilization (SS), and Synapse (Sy). (B) Bar graph showing the percentage of nodes within each biodomain graph that were identified as key drivers. The total number of nodes in each biodomain graph is shown parenthetically on the y-axis. (C) Processes driven by nodes in each biodomain graph. The difference in significance (-log10 FDR) from weighted versus unweighted key driver analyses is shown (y-axis) for processes within each biodomain (x-axis). (D) Network graph of biodomain interactions inferred from key driver analyses. The size of each node shows the number of nodes in each biodomain graph, the thickness of each edge shows the number of key drivers from the source domain graph for processes in the target domain. (E) Heatmap representation of the biodomain interactions inferred from key driver analyses. The number of key drivers in each graph is shown on the right, and the color shows the number of key drivers for processes in each biodomain along the x-axis from each graph along the y-axis. (F) The fraction of in-degree versus out-degree nodes for each biodomain in the interaction graph (D-E). The total number of key drivers identified in each biodomain graph are shown parenthetically along the y-axis.

**Table 1.** Top key drivers across biodomain networks by weighted FDR.

| Key Driver Node | Top Biodomain(s) | AMP-AD Nominated | Beckmann *et al* Key Driver | Sum ΔFDR | TREAT-AD TRS | TRS Percentile |
| --- | --- | --- | --- | --- | --- | --- |
| EGFR | Autophagy<br>Immune Response |  |  | 12.505 | 4.373 | 99.8% |
| FYN | Autophagy<br>Immune Response |  |  | 7.674 | 3.617 | 94.1% |
| CLTC | Endolysosome |  | TRUE | 14.448 | 3.171 | 83.6% |
| YWHAB | Endolysosome |  | TRUE | 3.839 | 3.729 | 95.7% |
| CTNNB1 | APP Metabolism |  |  | 5.027 | 3.423 | 90.2% |
| HRAS | Autophagy<br>Immune Response |  |  | 7.970 | 4.160 | 99.3% |
| DNM1 | Endolysosome<br>Synapse | TRUE | TRUE | 10.083 | 4.241 | 99.6% |
| CALM1 | Synapse |  |  | 9.754 | 3.313 | 87.4% |
| RAC1 | Autophagy<br>Immune Response |  |  | 8.928 | 4.109 | 99.1% |
| GRB2 | Autophagy<br>Immune Response |  |  | 8.864 | 3.326 | 87.7% |
| TP53 | Immune Response |  |  | 7.310 | 3.026 | 79.7% |
| SELENBP1 | APP Metabolism | TRUE |  | 3.870 | 3.799 | 96.7% |
| ALDH3A2 | Mitochondrial<br>Metabolism |  |  | 5.679 | 3.980 | 98.4% |
| DTNBP1 | Endolysosome<br>Synapse |  |  | 6.788 | 3.975 | 98.4% |
| RAB8A | Endolysosome<br>Synapse |  |  | 5.510 | 2.006 | 53.9% |

Although the biodomains have been useful in partitioning AD-relevant biology, many of these processes can and do interact within cells. Through mapping the cross-domain influences of key drivers we can build a more integrated model that goes beyond identifying drivers within a single domain and infers how perturbation in one cellular system could influence other cellular functions.

The terms impacted by each key driver can be assigned back to the parent biological domain in order to identify which domains are most strongly influenced by each graph (Figure 3C). Distinct sets of processes are driven by each graph. For example, the 35 key drivers identified in the Apoptosis graph are predicted to influence processes primarily from the Immune Response, Apoptosis, and Oxidative Stress domains, while the 26 key drivers identified in the Cell Cycle graph are predicted to influence primarily processes within the Mitochondrial Metabolism, Lipid Metabolism, and DNA Repair domains. These relationships — between hub nodes in the graph for one domain influencing processes in another domain — can be used to infer interactions between the biodomains (Figure 3D). Analyzing these interactions highlights the strong mutual influence between the Endolysosome and Synapse domains, the influence of Cell Cycle and Oxidative Stress on the Mitochondrial Metabolism domain, and the centrality of the Structural Stabilization domain. Visualizing the same interaction data as a heatmap provides additional detail around these domain level interactions (Figure 3E). For example, the graphs that contain the largest number of drivers for Synapse domain terms are Endolysosome, Structural Stabilization, and Proteostasis, whereas the graph containing the largest number of Structural Stabilization term drivers is the Structural Stabilization graph. Analyzing node degree in the inferred biodomain interaction graph reveals that the Metal Binding and Homeostasis, Structural Stabilization, and Myelination domains have the highest proportion of out-degree (Figure 3F). This suggests that these domains tend to influence processes in other domains more than they are being driven in the graphs of the other domains.

Our analyses highlight the importance of the Structural Stabilization, Endolysosome, and Lipid Metabolism domains, both through the fraction of nodes identified as key drivers (Figure 3B), and through the proportion of outgoing versus incoming inferred domain interactions (Figure 3F).

Focusing on cross-domain interactions, the Structural Stabilization graph contains the most key drivers for processes in the Synapse (29 drivers), Immune Response (22 drivers), Proteostasis (21 drivers), and Endolysosome (13 drivers) domains (Figure 3E). The key drivers within the Endolysosome graph are mostly for processes within the Synapse (41 drivers), Immune Response (20 drivers), Proteostasis (17 drivers), and Autophagy domains (6 drivers) (Figure 3E). Key drivers within the Lipid Metabolism domain are influencing processes concentrated within the Endolysosome (13 drivers), Immune Response (11 drivers), Structural Stabilization (11 drivers), and Synapse (10 drivers) domains (Figure 3E). Notably both the Synapse and Immune Response domains are among the domains influenced by each of these three most influential domains. Therefore the relationships between Structural Stabilization, Endolysosome, and Lipid Metabolism domain genes influencing functions relevant to Synapse and Immune Response domains seem to be key points of convergence between the AD biodomains, potentially suggesting how these domain associated processes interact to drive forward the neurodegenerative sequelae.

### 3.3 Integrating Key Drivers to Identify Genes Contributing to Disease Pathogenesis

There are several instances of distinct driver nodes identified for the same process across different biodomain graphs (Figure S5), likely due to the inclusion of different nodes into the respective graphs (Figure 2E). To identify which nodes are most consistently identified as a key driver of given processes, we integrated the key driver results across biodomain graphs (see Methods).

There 353 key drivers identified in more than one biodomain graph as top drivers, involving 169 unique driver nodes and 190 distinct processes (Supplemental Table 3). Several of the most consistently influential nodes are well supported by previous studies of AD and related biology.

Beta-Catenin (CTNNB1) is identified as a top key driver of the term “cellular response to amyloid-beta” (GO:1904646) from the APP Metabolism biological domain. CTNNB1 is identified as the top key driver of this process independently in six of the nine biodomain graphs (Figure 4A, Figure S5) and is identified as a hub for the 38 genes annotated to this term, despite not being annotated to the term itself (Figure 4B). CTNNB1 has an overall AD Target Risk Score among the top 10% of all scored targets (score 3.42, rank #2,353) and there is myriad evidence accumulated over the past 25 years of research that links beta-catenin and Wnt signaling, which acts upstream to promote stability of beta-catenin, with amyloid-beta pathology. There is evidence of altered Wnt/beta-catenin signaling in AD brains [27] as well as animal models [28]. Among the earliest lines of evidence supporting this association was the observation that PSEN1, involved in the proteolytic processing that produces amyloid-beta peptides, stabilizes beta-catenin and that AD-associated mutations reduce this stabilization [29]. Activation of Wnt signaling, on the other hand, can be protective against amyloid-beta induced neurotoxicity [30–32] and promote the blood-brain-barrier integrity [33], among other potentially beneficial effects. For these reasons Wnt/beta-catenin signaling has been proposed and is under active investigation as a therapeutic intervention in the treatment of AD [34–36].

**Figure 4.**
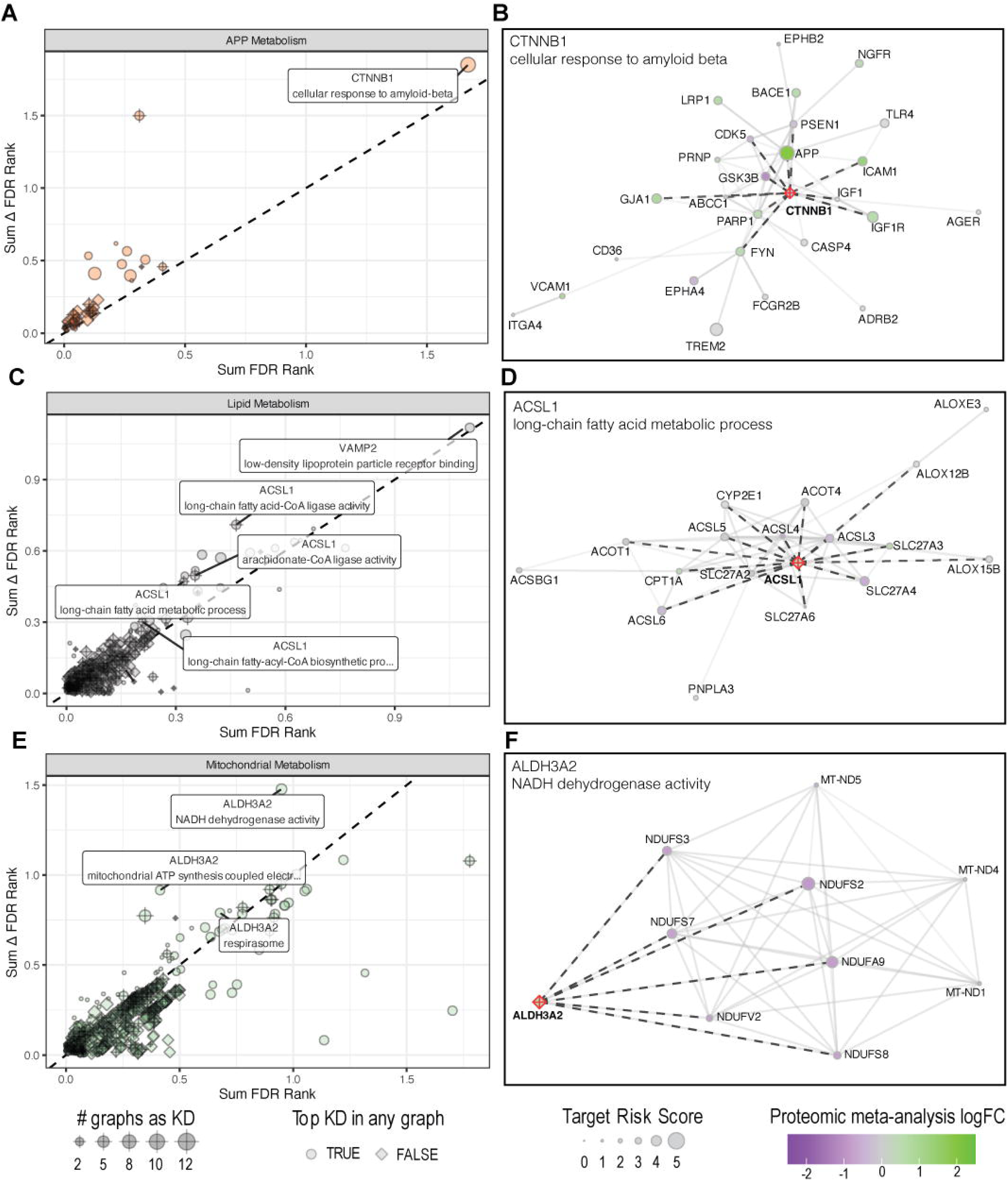
Example top reproducible key drivers across domains. The top key drivers for processes associated with APP Metabolism (A), Lipid Metabolism (C), and Mitochondrial Metabolism (E). Each plot shows the sum of the risk-weighted FDR rank (x-axis) versus the sum of the delta FDR rank (y-axis) for key drivers identified across multiple biodomain graphs. The size of each point corresponds to the number of biodomain graphs in which the node is found to be a driver for the identified process, and the shape of each point indicates whether the identified driver is the top driver for the specified term. Example key drivers are highlighted for each biodomain: CTNNB1 for “cellular response to amyloid beta” (B), ACSL1 for “long-chain fatty acid biosynthetic process” (D), and ALDH3A2 for “NADH dehydrogenase activity” (F). These examples show the key driver node (red) with the other proteins annotated to the process. Dashed lines indicate the shared edges between proteins in the process and the identified key driver. The size of the nodes corresponds to the Target Risk Score for each protein and the color represents the proteomic log fold change for the protein in AD patients versus control patients.

Another consistently identified top key driver within the biodomain graphs is the gene ACSL1, which is a key driver of both “long-chain fatty acid metabolic process” (GO:0001676) and “long-chain fatty acid-CoA ligase activity” (GO:0004467) terms, both of which are from the Lipid Metabolism biological domain. ACSL1 is the top key driver of these processes in two of the three biodomain graphs where it is identified as a hub (Figure 4C-D) and, unlike CTNNB1, ACSL1 is annotated to both terms for which it is identified as a key driver. ACSL1 has an overall AD Target Risk Score among the top 5% of all scored targets (score 3.72, rank #1,040) and has been nominated by AMP-AD investigators as a candidate therapeutic target in AD. ACSL1 has recently been associated with lipid-droplet accumulating microglial (LDAM) cells in AD [37]. While overexpression of ACSL1 had previously been reported to stimulate the formation of lipid droplets in other contexts [38,39], Haney *et al* demonstrated that a subset of microglia defined by the expression of ACSL1 are associated with a lipid-droplet phenotype. The abundance of ACSL1 expressing LDAM was highest in patients carrying two copies of the APOE ε4 allele, which is the strongest allele associated with AD risk from GWAS. Moreover, perturbation of ACSL1 in a CRISPR knock-out screen in iPSC derived microglial cells lead to an overall decrease in the lipid droplet phenotype in those cells. This supports the assertion from our key driver analyses that ACSL1, through regulation of fatty acid metabolism, could be a driver gene of disease-relevant processes.

Finally, ALDH3A2 is identified as a key driver of several processes from the Mitochondrial Metabolism biodomain relating to complex I of the electron transport chain — specifically “NADH dehydrogenase activity” (GO:0003954), “mitochondrial ATP synthesis coupled electron transport” (GO:0042775), and “respirasome” (GO:0070469). ALDH3A2 is the top key driver for these processes in all four biodomain graphs where it is identified as a hub (Figure 4E-F), but ALDH3A2 itself is not annotated to any of the terms for which it is identified as a key driver. ALDH3A2 has an overall Target Risk Score among the top 2% of all scored targets (score 3.98, rank #380), yet the only study to link ALDH3A2 with AD previously showed that rare variants in ALDH3A2 are associated with the risk of developing late-onset AD [40]. However, there are numerous studies that link the effects of combining non-specific inhibition of aldehyde dehydrogenases and oxidative phosphorylation on the growth of cancer cell lines. For example, there are synergistic effects of using non-specific inhibitors of aldehyde dehydrogenases and oxidative phosphorylation on chemotherapeutic effects on a variety of cancer cells [41], this is likely mediated through synergistic effects on mitochondrial functioning (e.g. ATP production) [42], and specific knock-down of ALDH3A2 reduced cell growth in leukemia cell lines [43]. Quantitative proteomic profiling of colorectal adenocarcinoma cell lines, sorted for high versus low aldehyde dehydrogenase activity, demonstrated that increased expression of aldehyde dehydrogenases is associated with decreased expression of several proteins involved in the mitochondrial electron transport chain as well as decreased expression of the NCSTN subunit of the gamma-secretase complex [44]. There is evidence that ALDH3A2 protein and mRNA is increased in expression in LOAD brains (Figure S6A) and this expression seems to be most pronounced among glutamatergic neurons (Figure S6C). Data from iPSC-derived glutamatergic neurons from CRISPRbrain.org suggest that increasing the expression of ALDH3A2 is associated with a significantly decreased survival of neurons in culture, although it does not meet the criteria to classify as a hit in this assay (Figure S6B). These results along with the observations from cancer cell lines propose a model wherein increased expression of ALDH3A2 is associated with relative decreases in oxidative phosphorylation - a prominent feature from the analysis of post-mortem AD brains - and that this may contribute to the loss of glutamatergic neurons in disease.

## 4. DISCUSSION

We developed an approach to reconstruct AD biodomain-specific graphical networks around the most disease associated proteins from each domain and then used these graphs to identify key driver nodes. We started with a composite interaction graph from Pathway Commons [15], a superior approach for disease gene identification [9,10]. We applied stringent filters for brain-expressed nodes and literature-supported edges, which reduced the overall network size by removing 43.8% of nodes and 78.7% of edges from the full graph. Biodomain graphs were generated starting with the most risk- enriched nodes (i.e. leading edge genes) from each biodomain. We then used Dijkstra’s algorithm for graph reconstruction using the shortest paths between nodes, focusing on high-scoring, domain- relevant nodes to build out the graphs. This process enumerated the interactions between disease risk genes and resulted in the generation of biodomain-specific molecular interaction graphs with limited overlap between domains. We identified influential hub nodes (key drivers) within these graphs using wKDA, prioritizing drivers based on AD risk weights. The identified key drivers are enriched for previously identified AD drivers [14], high-priority AMP-AD nominated genes, with a suggestive enrichment of GWAS loci. Our analysis integrates more data sources than previous KDA studies [14] and associates key drivers to specific AD functions, allowing us to predict their role in disease. Placing drivers into the context of a biological pathway (or intersecting set of pathways) generates specific hypotheses about their relationship to AD pathogenesis.

We note that there are multiple key driver nodes prioritized for the same process based on the biodomain graph context (Figure S5). This is expected, as the biodomain graphs are only partially overlapping and certain nodes and sub-networks will be differentially present across graphs. We therefore focused on key drivers consistently identified across multiple networks, as their influence is robust to changing sub-network contexts. Integrating results across biodomains highlights nodes with process-specific disease influence. Three high priority key drivers emerged from our analyses. First, CTNNB1 is a key driver for “cellular response to amyloid-beta” within the APP metabolism domain (Figure 4A-B), a role supported by two decades of research [29,34]. Second, ACSL1 influences Lipid Metabolism in AD, a finding supported by more recent work showing ACSL1 regulates lipid droplet formation in microglia and was found to be a marker of lipid droplet accumulating microglia in AD brains [37]. This suggests a strong modifier role in innate immunity and links lipid droplet formation to disease risk. Third, ALDH3A2 is a key driver for Mitochondrial Metabolism, directly linking to mitochondrial risk genes within the NDUFS family, such as NDUFS2 (Figure 4E-F). This demonstrates a pathway-level association between ALDH3A2 and oxidative phosphorylation initiation through complex I of the electron transport chain. This is notable because mitochondrial complex I function was one of the strongest disease risk signatures in our previous work [4]. While there is no literature linking the influence of aldehyde dehydrogenases on oxidative phosphorylation in AD, there is evidence of their interdependence from the cancer literature [41], supporting the hypothesis that ALDH3A2 may directly modulate the ETC in AD pathogenesis as well. These examples are non-exhaustive, as over 160 other consistent key drivers warrant further study as candidate AD therapeutic targets. Experimentally validated nodes will undergo further resource development within the TREAT-AD consortium and be made publicly available.

We also used wKDA results to infer regulatory relationships between biodomains. Although biodomains silo gene function into distinct molecular processes, these processes do not act in isolation and likely interact to influence disease pathogenesis. A continuing goal when partitioning disease risk into biological domains has been to identify these points of convergence and to understand how these interactions can influence disease progression. Our analyses highlight the influence of the Structural Stabilization, Endolysosome, and Lipid Metabolism domains on processes from both the Synapse and Immune Response domains. Notably, a recent graph neural network machine learning framework, which leverages the AD biological domains, also highlighted the central influence of the Lipid Metabolism and Endolysosome domains [13]. The top processes targeted in our cross-domain interactions involve synaptic vesicle endocytosis and transport (Synapse domain), and phagocytosis and neuroinflammatory response (Immune Response domain). These interfaces are interesting to consider as potential therapeutic targets because they represent broader, systems-level processes that could impact disease pathogenesis through multiple relevant mechanisms.

Our approach has several limitations. First, it relies on existing annotations, including gene-to- GO term mappings and Pathway Commons interactions; poorly annotated genes and interactions will be missed. For example, 2,916 brain-expressed genes are absent from the Pathway Commons database and therefore will not be included in our models. Second, top key driver nodes may not be ideal therapeutic targets. Experimental validation is needed to confirm these predicted associations and the hypotheses implied by the topology of the key driver placement within the biodomain networks. However, the graph context allows exploration of adjacent nodes to identify targets that are more efficacious or less toxic than the top identified driver. The current implementation also treats all interactions equally, though different interaction types (e.g. regulatory) may be more informative than others (e.g. binding). Interactions are also scored independently of cell-type context, though the two proteins may be expressed in distinct cell types or across cellular boundaries. Future development will include building cell type specific graphs to investigate how biodomains operate within and between different cell types; for example, understanding Immune Response drivers in astrocytes versus microglia. Ultimately our approach is a tool that links driver genes to disease-implicated biology. This approach is an amalgamation of our endophenotypic mapping, disease risk modeling, and network analyses to accelerate target nomination. High priority targets with plausible hypothetical mechanisms of action that arise from these analyses warrant experimental validation in brain relevant cell types and models.

### 4.1 Conclusion

This work extends the tools and resources we developed previously [4], which represented the largest integration of genetic and multi-omic signatures of AD risk along with a system to localize signals of risk into specific disease molecular endophenotypes. Here, we contextualized those domains of risk into protein interaction networks and identification of the top risk implicated hub proteins as key drivers of disease relevant processes. Our approach identifies robust and consistent drivers across networks, highlighting both known AD-related proteins and novel candidates. A specific advantage of our approach is the interpretability. For example, we link the driver CTNNB1 to cellular responses to amyloid beta, a role which is well-supported by the literature. Other key drivers are identified along with their predicted impact on specific processes, suggesting numerous testable hypotheses. Our approach also enables inference of cross-domain influences, highlighting the strong impact of the Structural Stabilization, Endolysosome, and Lipid Metabolism domains across other domains. Mapping these interactions enables the consideration of systems-level interventions — proteins and processes that when perturbed may have larger impacts on disease relevant biology as they are predicted to influence multiple domains in concert. These domain interactions may also help identify candidate combination therapies given the multifarious manifestation of AD. Ultimately the Emory-Sage-SGC-JAX TREAT-AD center will use the results from these and other analyses to promote specific dark targets for future Target Enabling Package (TEP) development programs.

Resources developed from these TEPs, including informatic packages and experimental tools, are openly available to the scientific community. These TEPs will support therapeutic development for understudied targets that are currently intractable due to insufficient or ineffective tools and resources.

## ACKNOWLEDGEMENTS

The authors would like to acknowledge Dr. Robert Butler III for his insightful feedback during the revision of this manuscript. Data used in this study were obtained from the Accelerating Medicines Partnership Program for Alzheimer’s Disease (AMP AD) Consortium members below: Mayo RNAseq Study: Study data were provided by the following sources: The Mayo Clinic Alzheimer’s Disease Genetic Studies, led by Dr. Nilufer Ertekin Taner and Dr. Steven G. Younkin, Mayo Clinic, Jacksonville, FL, using samples from the Mayo Clinic Study of Aging, the Mayo Clinic Alzheimer’s Disease Research Center, and the Mayo Clinic Brain Bank. Data collection was supported through funding by NIA grants P50 AG016574, R01 AG032990, U01 AG046139, R01 AG018023, U01 AG006576, U01 AG006786, R01 AG025711, R01 AG017216, R01 AG003949, NINDS grant R01 NS080820, CurePSP Foundation, and support from Mayo Foundation. Study data include samples collected through the Sun Health Research Institute Brain and Body Donation Program of Sun City, Arizona. The Brain and Body Donation Program is supported by the National Institute of Neurological Disorders and Stroke (U24 NS072026 National Brain and Tissue Resource for Parkinson’s Disease and Related Disorders), the NIA (P30 AG19610 Arizona Alzheimer’s Disease Core Center), the Arizona Department of Health Services (contract 211002, Arizona Alzheimer’s Research Center), the Arizona Biomedical Research Commission (contracts 4001, 0011, 05 901, and 1001 to the Arizona Parkinson’s Disease Consortium), and the Michael J. Fox Foundation for Parkinson’s Research. Religious Orders Study/Memory and Aging Project (ROSMAP): We are grateful to the participants in the Religious Order Study and the Memory and Aging Project. This work was supported by the US National Institutes of Health (U01 AG046152, R01 AG043617, R01 AG042210, R01 AG036042, R01 AG036836, R01 AG032990, R01 AG18023, RC2 AG036547, P50 AG016574, U01 ES017155, KL2 RR024151, K25 AG041906 01, R01 AG30146, P30 AG10161, R01 AG17917, R01 AG15819, K08 AG034290, P30 AG10161, and R01 AG11101). Mount Sinai Brain Bank (MSBB): This work was supported by grants R01AG046170, RF1AG054014, RF1AG057440, and R01AG057907 from the NIH/NIA. R01AG046170 is a component of the AMP AD Target Discovery and Preclinical Validation Project. Brain tissue collection and characterization was supported by NIH HHSN271201300031C.

## CONSENT STATEMENT

No human participants were recruited for this work. The details of the Institutional Review Board (IRB)/oversight body that provided approval or exemption for the research described are given as follows: Western Institutional Review Board—Copernicus Group (WCG) IRB of Sage Bionetworks gave ethical approval for this work.

