## Supplemental Figures for "Mapping cross-domain drivers of Alzheimer’s disease risk through integrated network analysis"

Manuscript

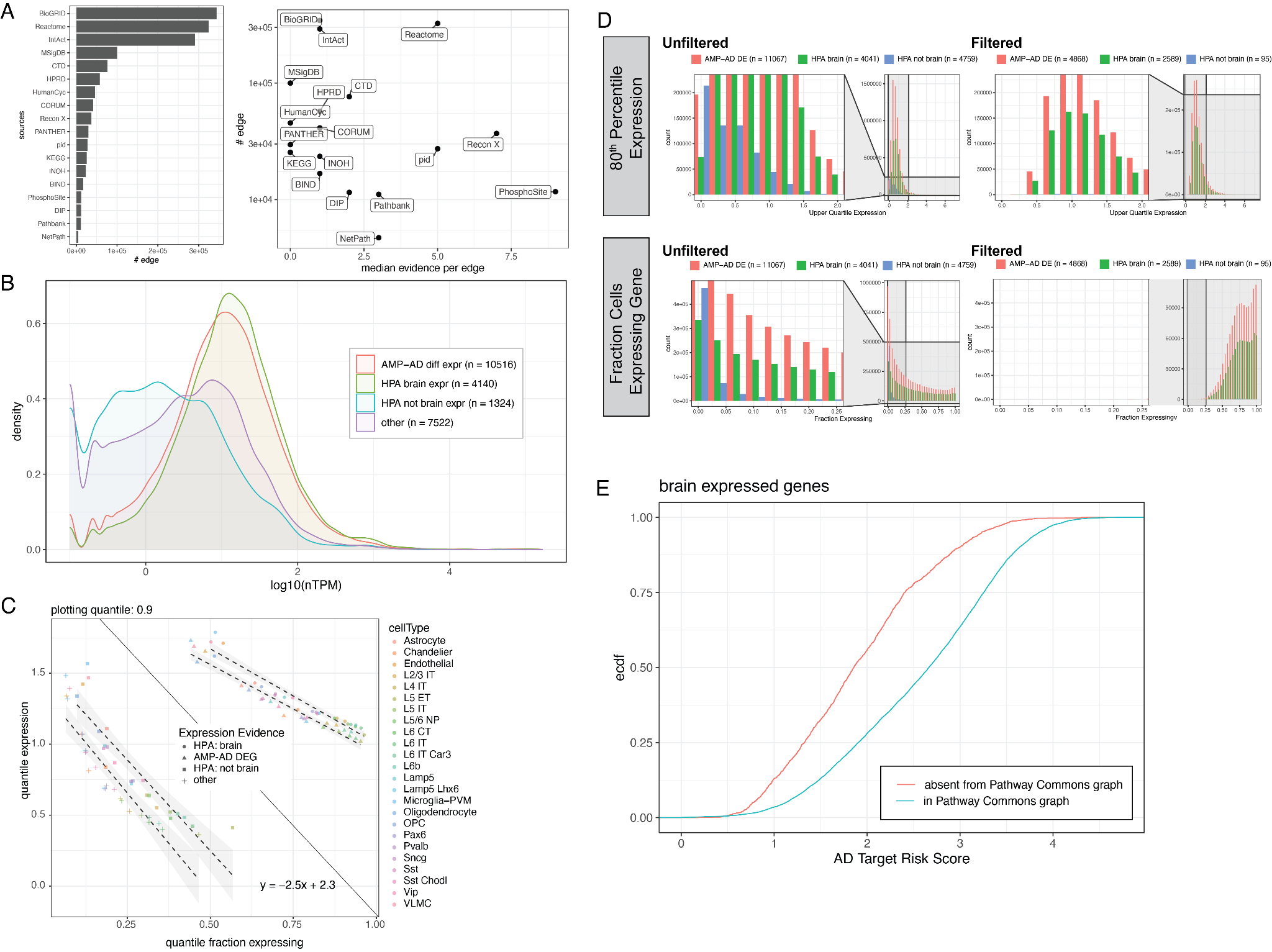

**Figure S1. Pathway Commons network filtering.** (A) Number of edges contributed by each data source to the Pathway Commons network and the median number of papers supporting each edge from the literature. (B) Log10 normalized TPM expression in GTEx brain for genes categorized by AMP-AD differential expression, Human Protein Atlas brain expression, or other. (C) Summarized single-cell RNA-seq data from SEA-AD showing the fraction of cells expressing genes versus their summary expression level per cell type, categorized by expression evidence class. (D) Abundance of genes in different expression classes before and after filtering the SEA-AD dataset, shown by fraction of cells expressing and upper quartile expression. (E) Cumulative distribution of AD Target Risk Scores for brain-expressed genes, comparing those present in the Pathway Commons graph versus those absent.

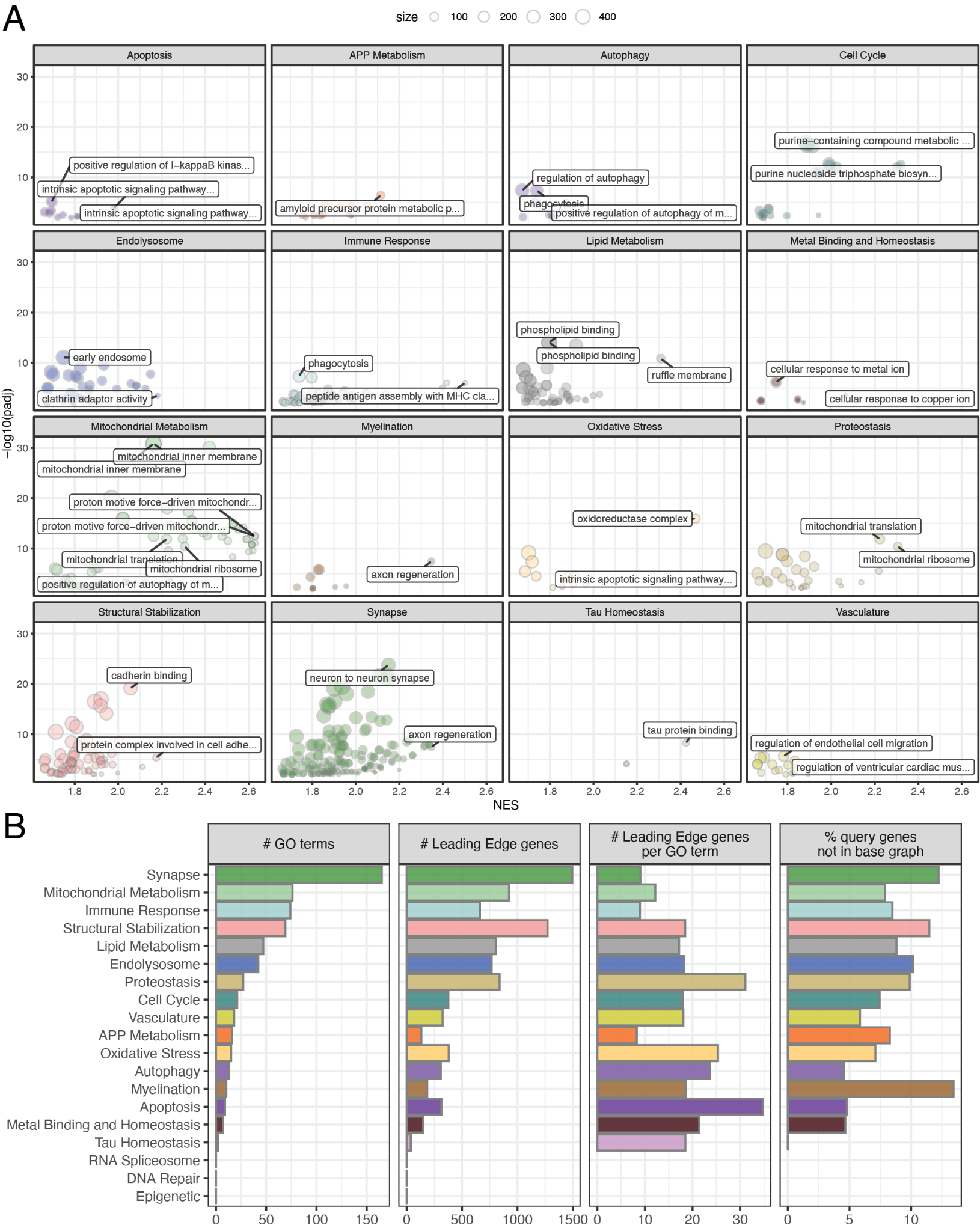

**Figure S2. Risk-enriched GO terms and leading edge genes by biodomain.** (A) Plots showing GSEA-enriched Gene Ontology (GO) terms for different biodomains. The x-axis represents the Normalized Enrichment Score (NES), and the y-axis represents the statistical significance (-log10(padj)). (B) Summarized characteristics of GO terms and leading edge genes across different biodomains.

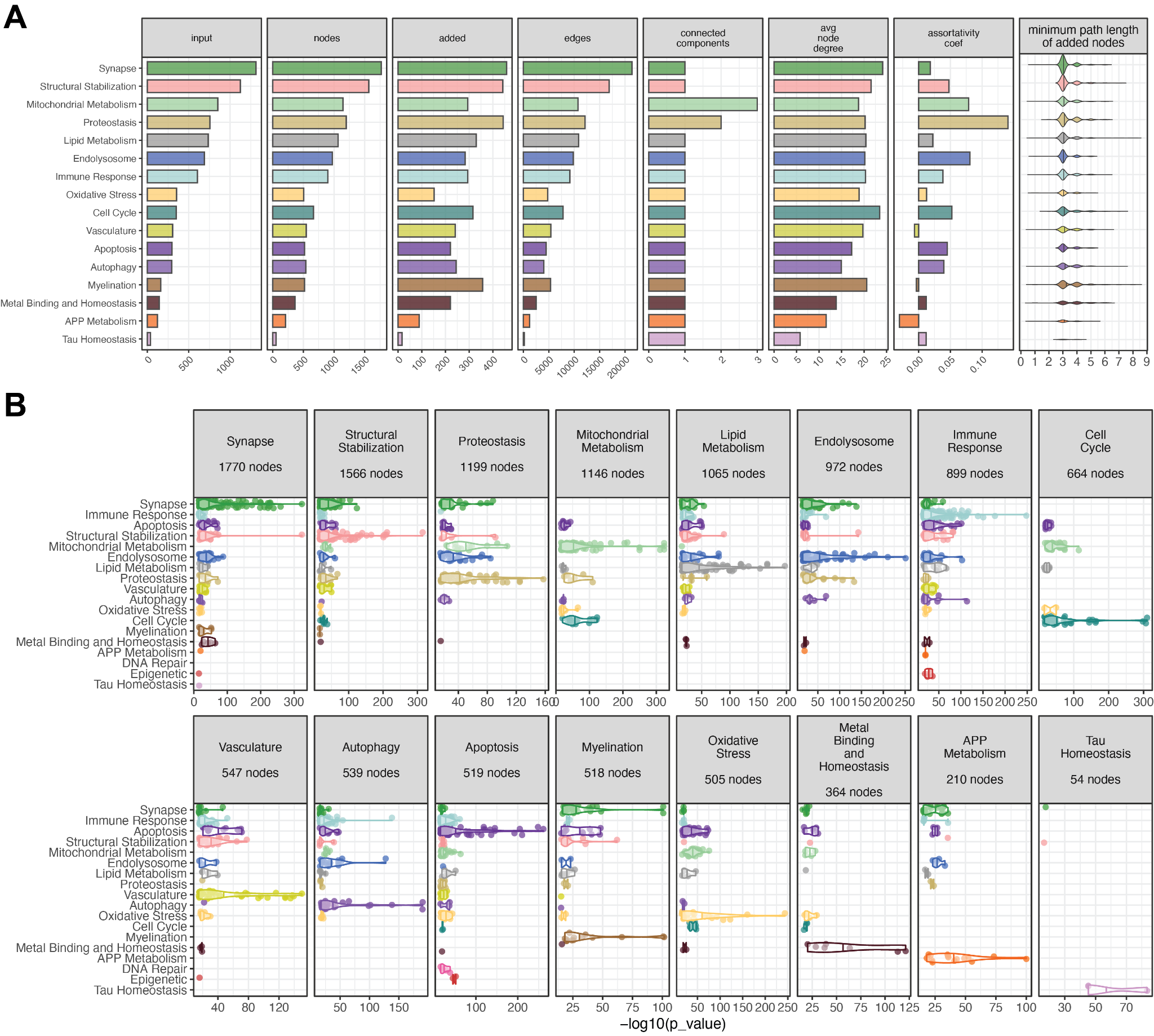

**Figure S3. Graph properties for biodomain networks.** (A) Network properties of AD biodomain networks including: number of input nodes, number of total nodes in the final graph, number of added nodes, number of edges, number of connected components, average node degree, assortativity coefficient, and the minimum path length of nodes added during tracing. (B) GO term over-representation analysis results mapped to AD biodomains for each biodomain network. For each network, the total number of nodes used for ORA is shown in the header.

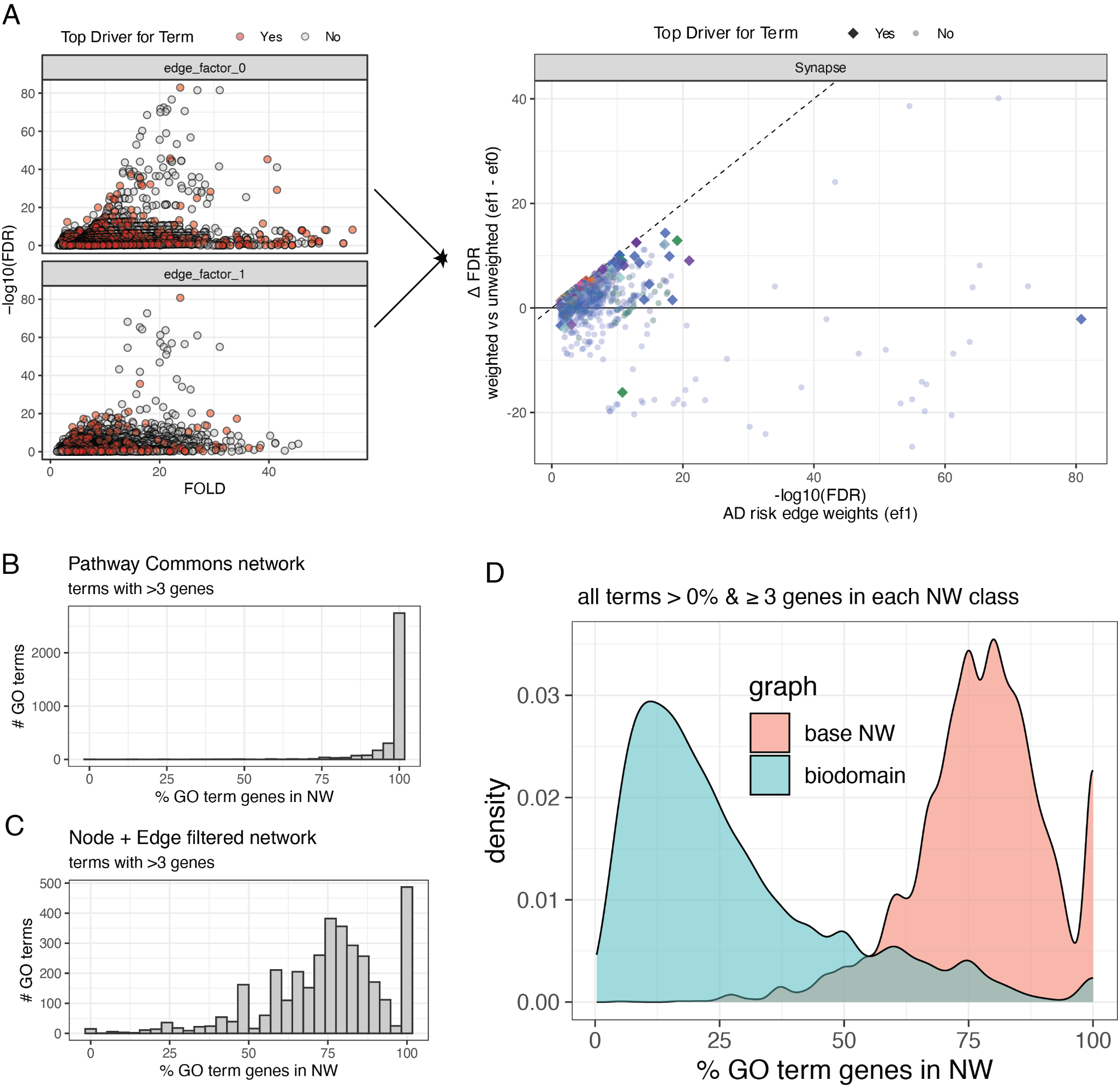

**Figure S4. Representation of GO terms in the key driver analysis results.** (A) Example analysis workflow to integrate the risk weighted KDA (“edge_factor_1”) with the unweighted KDA (“edge_factor_0”). The example results for the Synapse domain network are shown (left). The two runs were then integrated using the identified driver node and biodomain term and the difference between the significance in the weighted versus unweighted analyses (Δ FDR, y-axis) is plotted against the significance in the weighted analysis (x-axis). For each key driver shown, the color of the point reflects the parent domain of the process being driven. (B) Distribution of percent of genes from a GO term (with more than 3 genes) present in the full Pathway Commons network. (C) Similar to panel B, but for the node and edge filtered base network used as the starting point for biodomain graph generation. (D) Density plot of GO term gene representation in the base network (similar to panel C) versus the final biodomain specific networks. Most GO terms have very low representation of constituent genes within the biodomain networks (< 25%).

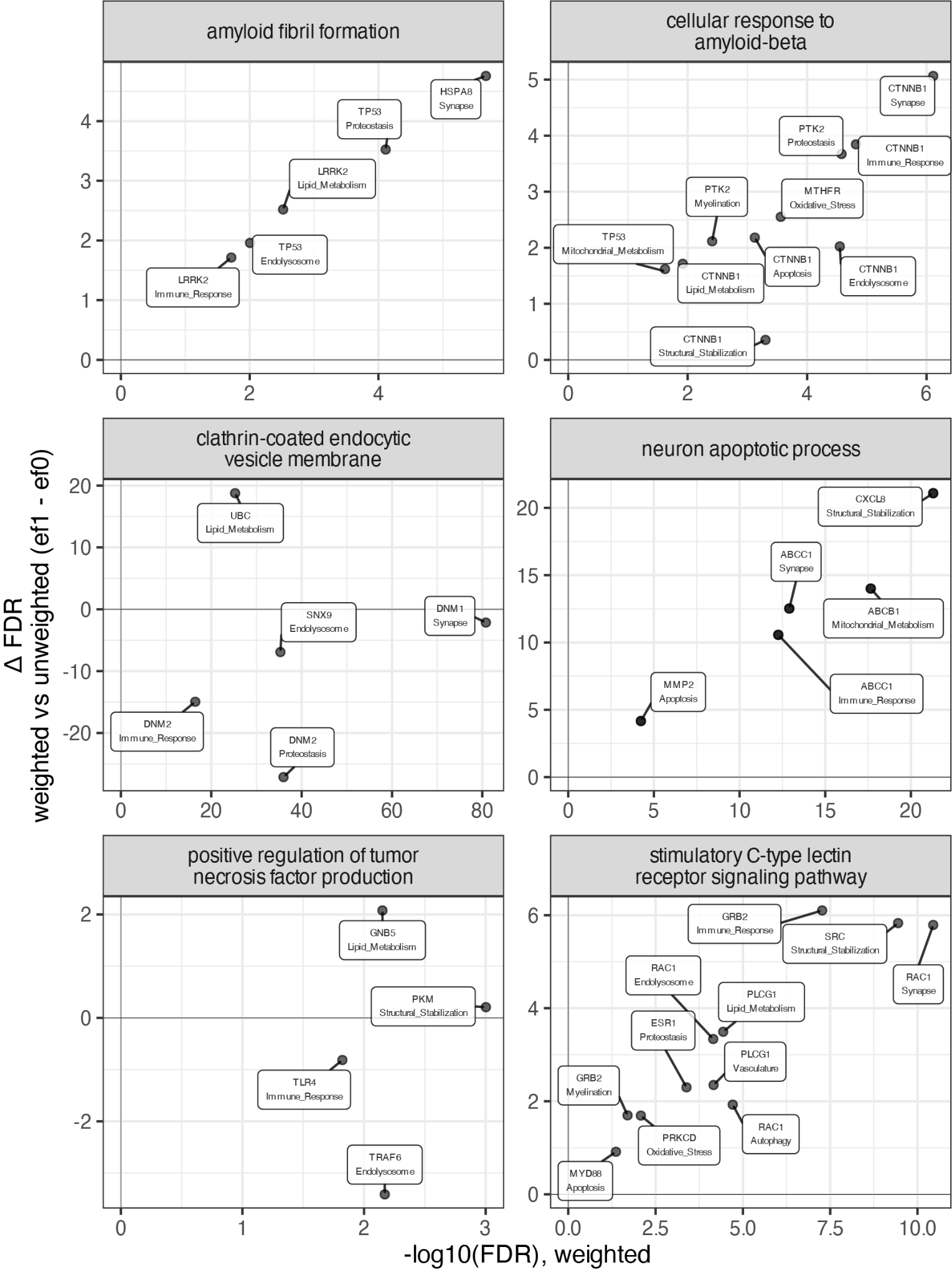

**Figure S5. Biodomain processes with distinct drivers across networks.** Key drivers identified for six terms are shown. The x-axis is the weighted -log10 FDR for the identified key driver and the y-axis shows the Δ FDR between unweighted and weighted analyses. Each point is labelled with the identity of the key driver node and the biodomain network in which it was identified.

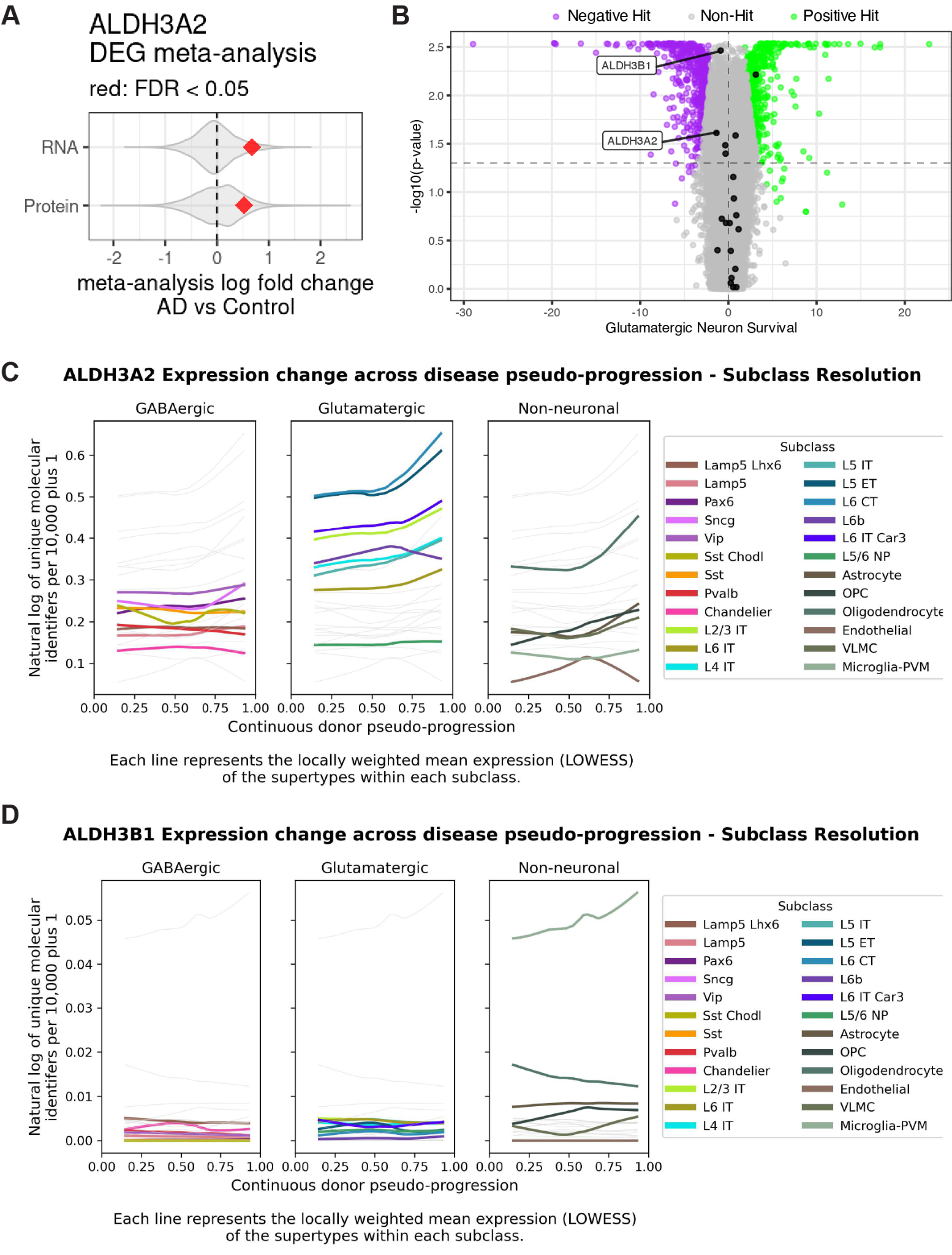

**Figure S6. ALDH3A2 expression patterns and functional genomics.** (A) Violin plots showing the distribution of all log fold change values for RNA and proteins calculated from the meta-analysis of AMP-AD datasets from Cary et al 2024. The red point shows the log fold change value of ALDH3A2 that is significantly up in both AMP-AD transcriptomics and proteomics. (B) Volcano plots of data from the CRISPRbrain resource showing the results of CRISPRa screening of genes influencing glutamatergic neuronal survival. Genes that increase survival when up-regulated are on the right of the x-axis, while genes that lead to a decrease in neuronal survival when up-regulated are on the left of the x-axis. Black points represent aldehyde dehydrogenase genes (“ALDH”), and the top two that negatively impact neuronal survival are shown (ALDH3B1 and ALDH3A2). Line plots showing the expression level for ALDH3A2 (C) and ALDH3B1 (D) across continuous donor pseudo-progression from the SEA-AD resource. Each line represents the locally weighted mean expression values of the supertypes within each subclass.
